# Stronger premicrosaccadic sensitivity enhancement for dark contrasts in the primate superior colliculus

**DOI:** 10.1101/2024.09.08.611886

**Authors:** Wenbin Wu, Ziad M. Hafed

**Author notes:** Correspondence: Ziad M. Hafed, Werner Reichardt Centre for Integrative Neuroscience, Otfried-Müller Str. 25, Tübingen, Germany, 72076.

## Abstract

Microsaccades are associated with enhanced visual perception and neural sensitivity right before their onset, and this has implications for interpreting experiments involving the covert allocation of peripheral spatial attention. However, the detailed properties of premicrosaccadic enhancement are not fully known. Here we investigated how such enhancement in the superior colliculus depends on luminance polarity. Rhesus macaque monkeys fixated a small spot while we presented either dark or bright image patches of different contrasts within the recorded neurons’ response fields. Besides replicating premicrosaccadic enhancement of visual sensitivity, we observed stronger enhancement for dark contrasts. This was especially true at moderate contrast levels (such as 10-20%), and it occurred independent of an individual neuron’s preference for either darks or brights. On the other hand, postmicrosaccadic visual sensitivity suppression was similar for either luminance polarity. Our results reveal an intriguing asymmetry in the properties of perimicrosaccadic modulations of superior colliculus visual neural sensitivity.

## Introduction

Microsaccades are small saccades that occur during periods of maintained gaze fixation ^1–4^. The generation of these rapid eye movements is associated with strong motor bursts in the superior colliculus (SC) ^2,5–7^, and these motor bursts are similar in character to those associated with larger saccades ^8^. As a result, microsaccades are additionally known to be accompanied by robust perimovement alterations in visual neural sensitivity and perception, just like larger saccades are ^2,7,9^.

Perimicrosaccadic alterations in SC visual responses include premicrosaccadic enhancement of neural sensitivity ^10^, postmicrosaccadic suppression of neural sensitivity ^11–13^, as well as perimovement shifts in foveal visual response field (RF) locations ^7,14^. These alterations are consistent with known perimicrosaccadic changes in perception ^9,15,16^. For example, premicrosaccadic enhancement in perception occurs for peripheral stimuli, and this also happens in covert attentional cueing tasks ^15^. This suggests that perimicrosaccadic alterations in visual processing (whether in the SC, other brain areas, or perceptually) can have a substantial impact ^9^ on the interpretation of experiments containing sensory transients, such as in the case of cueing paradigms for the study of covert attention ^9,15–17^. In fact, coupled with the intrinsic rhythmicity of microsaccades and the resetting of such rhythmicity by exogenous peripheral or foveal visual stimuli ^9,18,19^, it is possible ^9,15,16,20^ (at least theoretically) to fully account for classic Posner-style ^21–23^ cueing effects solely by the known perimicrosaccadic alterations in vision. This makes it necessary to keep studying additional properties of perimicrosaccadic effects.

Intriguingly, perisaccadic changes in visual sensitivity can exhibit different asymmetries as a function of different visual scene parameters. For example, even though visual perception can be worse in the upper rather than lower retinotopic visual field in the absence of saccades ^24–31^, perisaccadic perceptual performance is actually better in the upper visual field (during the strong perisaccadic suppression epoch) ^13^; this is an opposite asymmetry from the no-saccade case, but it is one that nonetheless matches the one present in SC visual neural sensitivity ^32^. Moreover, the strength of perceptual suppression associated with microsaccades can vary as a function of the visual appearance of foveated images ^33,34^, and different image properties can modulate the strength of perisaccadic suppression both perceptually and neurally ^12,34–37^.

Here we uncovered an additional asymmetry in perimicrosaccadic visual sensitivity, this time with a focus on SC visual responses to different luminance polarities. We previously found that SC visual responses can be stronger for either dark or bright luminance polarities, and that such visual responses still occur slightly earlier for dark stimuli, regardless of whether individual neurons prefer (in terms of their visual response strengths) darks or brights ^38^. We also discussed this in light of known asymmetries in the visual processing of darks and brights ^39–41^. For example, dark contrasts in natural scenes tend to co-occur with regions of low spatial frequency image content and “far” binocular disparities ^42,43^. This suggests that foveating eye movements, which are mediated by the SC, can have shorter reaction times for dark than bright stimuli. This is indeed the case ^38^, adding further evidence ^44^ that the SC’s visual tuning preferences for low spatial frequencies ^45^ and for image regions dominated by far binocular disparities ^32^ can have a direct influence on visually-guided eye movement behavior. Importantly, it was not previously studied how visual response strength for dark and bright stimuli was altered by microsaccades. This is what we did here.

We found a specific asymmetry in the premicrosaccadic interval: premicrosaccadic enhancement of peripheral SC visual neural sensitivity was stronger for dark than bright stimuli, but only at moderate contrast levels, and this was independent of whether individual neurons intrinsically preferred one or the other luminance polarity. Postmicrosaccadic suppression of visual neural sensitivity was similar for either luminance polarity. These results add to the growing evidence of asymmetries in visual processing, and they extend them to the case involving concomitant eye movement generation. These results also suggest that a stronger premicrosaccadic enhancement for dark stimuli of moderate contrasts can help equalize SC contrast sensitivity curves, bringing them nearer to their saturation regime for a wider range of contrast levels. In other words, because SC neuronal contrast sensitivity curves are characterized by a monotonic increase in firing rates followed by a saturation phase at higher contrast levels ^10,38,46^, increasing the sensitivity of neurons for only the moderate contrasts implies that the neurons reach their saturation regime for a larger range of possible stimulus energies. This allows for enhanced premicrosaccadic detection of stimuli in which dark contrasts might be particularly prevalent in natural scenes ^42^, and it is a phenomenon that very likely also happens before larger saccadic eye movements as well.

## Results

We investigated how the visual response strength of extrafoveal SC neurons is altered if the visual stimulus appearing within the neurons’ RF’s occurs before or after tiny microsaccades. To do so, we trained our monkeys to fixate a small spot, and we presented a disc of 0.51 deg radius within extrafoveal SC RF’s (Methods). Across trials, the disc could be brighter than the gray display background (positive luminance polarity) or it could be darker (negative luminance polarity). We varied the Weber contrast of the disc (relative to the background luminance) from trial to trial (Methods). In post-hoc analyses, we separated trials according to when the stimulus onset occurred relative to a given microsaccade (Methods). Baseline trials were those in which there were no microsaccades occurring within less than +/- 150 ms from stimulus onset. This was a sufficiently long interval to avoid at least the largest modulatory effects of microsaccades on vision ^47^. For a given stimulus luminance polarity and contrast, we then measured the peak visual response in baseline trials, and we related it to trials in which the stimulus onset occurred within different time epochs around microsaccade onset.

### Stronger premicrosaccadic enhancement for moderate dark contrasts

Figure 1a-d shows the visual responses of an example SC neuron to two different contrasts (Fig. 1a, b versus Fig. 1c, d) and two different luminance polarities (Fig. 1a, c versus Fig. 1b, d). The dark and light gray curves show the baseline firing rates of the neuron within each luminance polarity condition in the complete absence of microsaccades near stimulus onset (Methods). This example neuron preferred the dark luminance polarity, as evidenced by the stronger visual responses for the negative luminance polarity at both shown contrast levels (compare the dark and light gray curves). The curves with the different shades of blue in the figure show the same neuron’s visual response strengths, but now when the stimulus onsets occurred within <100 ms before microsaccade onset. The microsaccades had an average amplitude of only approximately 0.45 deg, whereas the neuron preferred an extrafoveal eccentricity of approximately 6.3 deg. For the dark stimuli (negative luminance polarity), premicrosaccadic enhancement in visual sensitivity of the neuron was evident (Fig. 1b, d).

**Figure 1.**
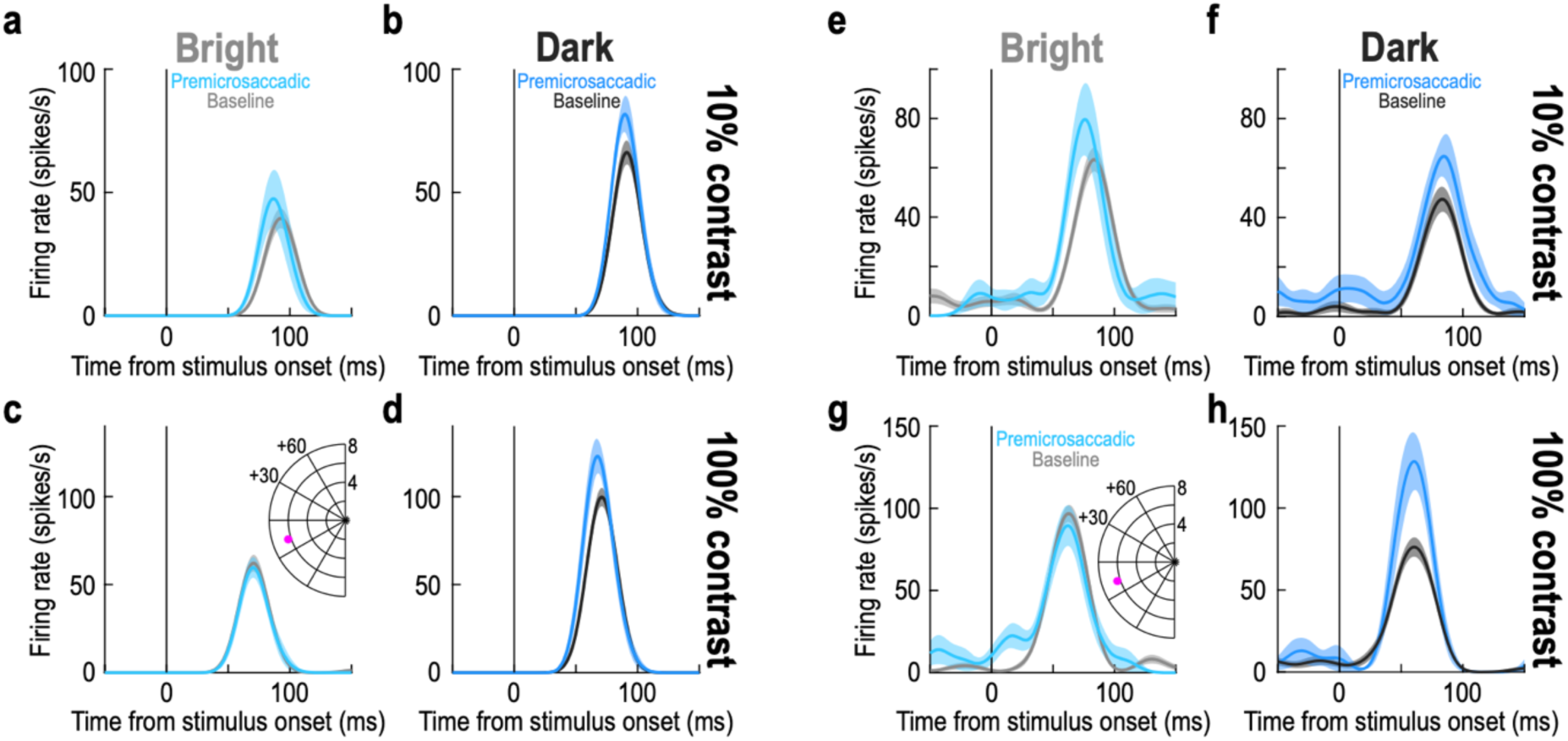
Stronger premicrosaccadic enhancement of superior colliculus (SC) visual sensitivity for dark contrasts. **(a, b)** Visual responses of an example SC neuron to 10% contrast stimuli of either positive (**a**) or negative (**b**) luminance polarity. The gray curves (dark in **b** and lighter shade in **a**) show the stimulus-evoked visual responses of the neuron in the absence of any microsaccades within +/- 150 ms from stimulus onset. The neuron preferred the dark stimulus. The colored curves show the same neuron’s visual responses when stimulus onset occurred within <100 ms before microsaccade onset. Premicrosaccadic enhancement occurred, consistent with our earlier results ^10^, but it was stronger for the dark contrast. **(c, d)** The same neuron’s responses for 100% contrast stimuli. This time, there was no premicrosaccadic enhancement for bright stimuli (c), but it was present for dark stimuli (**d**). The inset shows the retinotopic preferred RF location of the neuron (approximately 6 deg eccentricity in the lower left visual field). **(e-h)** Another example neuron from the same SC site. This neuron preferred bright stimuli in its baseline curves. Once again, it exhibited premicrosaccadic enhancement. However, the enhancement was weak or absent for bright stimuli, and more robust for dark stimuli (e.g. the 100% contrast condition). Thus, there was stronger premicrosaccadic enhancement of SC visual sensitivity for negative luminance polarity stimuli. Error bars denote SEM. n = 24-33 trials in the baseline curves across panels; n = 5-9 trials in the premicrosaccadic curves across panels (Methods).

On the other hand, for the bright stimuli (positive luminance polarity), the enhancement was weak or absent (Fig. 1a, c). Similar observations were also made for a second example neuron from the same monkey; this time, the neuron (which was recorded simultaneously with the other neuron) preferred bright rather than dark stimuli in the absence of microsaccades (Fig. 1e-h, gray baseline curves). Once again, the neuron exhibited robust premicrosaccadic enhancement of visual sensitivity, and particularly for the negative luminance polarity (see, for example, the 100% contrast condition for this neuron; we later elaborate on the quantitative enhancement strengths as a function of different contrasts, luminance polarities, and neuron preferences for darks and brights). Therefore, regardless of neural preference for either darks or brights, premicrosaccadic enhancement was present in both of these example SC neurons, consistent with our earlier findings ^10^. More importantly, premicrosaccadic enhancement seemed to be more robust for the dark stimuli in these two neurons. We next inspected these observations in more detail across the entire population.

Across the population, we first confirmed that we obtained consistent premicrosaccadic enhancement ^10^ (and postmicrosaccadic suppression ^10–12^). To do so, we used an approach similar to one we used previously ^11^. We normalized each neuron’s firing rate curves across all trials, before pooling across neurons. The normalization factor was the peak of the average firing rate curve of the condition evoking the highest response in the absence of peristimulus microsaccades (Methods). Then, for each time interval in which a stimulus onset occurred relative to a given microsaccade onset, we plotted the average normalized peak visual response across all data. This resulted in time courses of perimicrosaccadic modulations of visual neural sensitivity for all stimulus conditions, which we summarize in Fig. 2. In this figure, each horizontal dashed line shows the average normalized response of the population in the baseline situation of no microsaccades within +/- 150 ms from stimulus onset (Methods). The correspondingly-colored solid curve shows the same response as a function of when a stimulus appeared relative to a given microsaccade onset (each sample on the x-axis shows an averaging window of within +/- 25 ms from the sample center). As can be seen, we systematically observed premicrosaccadic enhancement and postmicrosaccadic suppression of SC visual sensitivity, consistent with our earlier observations ^10–12^, and this was additionally the case for different contrasts and different luminance polarities.

**Figure 2.**
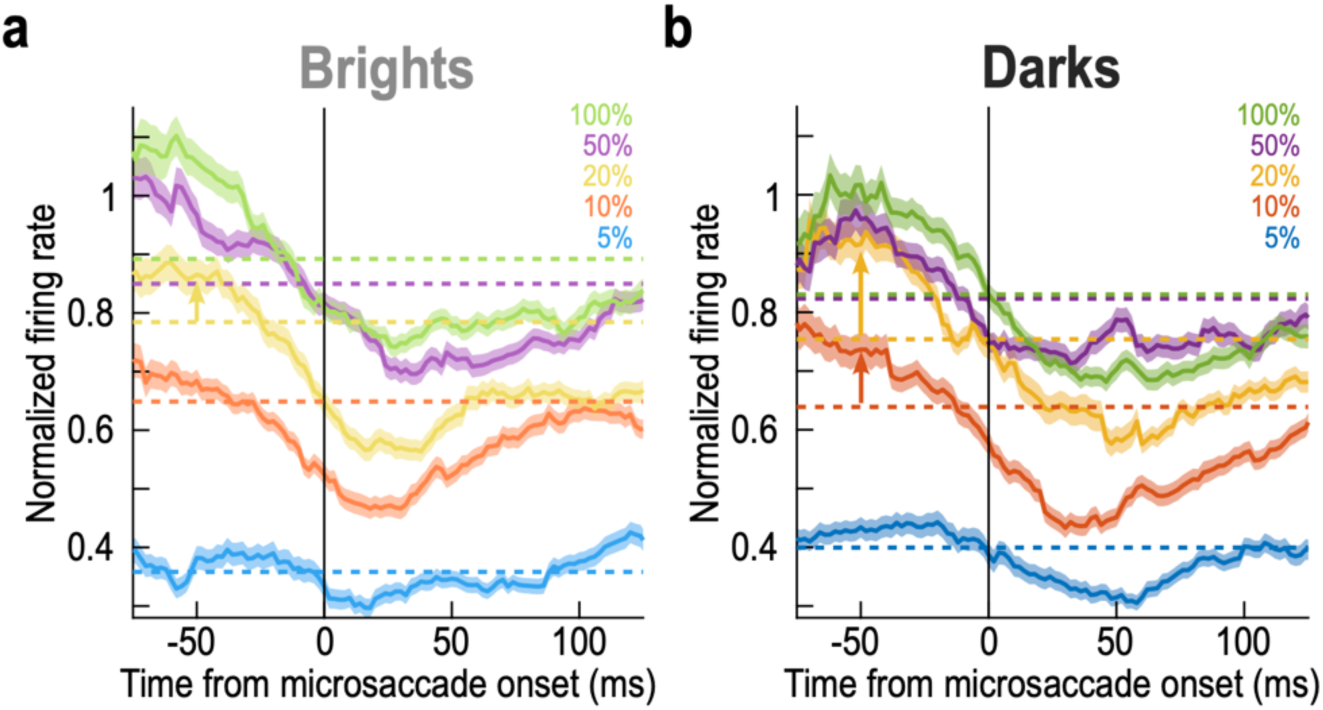
Stronger premicrosaccadic response gain enhancement for moderate dark contrasts in the primate SC. **(a)** For positive luminance polarity stimuli (brights), each dashed horizontal line shows the average normalized population firing rate (visual response strength) for trials in which stimuli of a given contrast appeared in the absence of any microsaccades within +/- 150 ms from stimulus onset (baseline trials). For each correspondingly-colored solid curve, the x-axis value indicates the center of a measurement window of +/- 25 ms from the given value, and the y-axis value indicates the normalized population visual response if stimulus onset occurred within the specific time window relative to microsaccade onset indicated by the x-axis value. For example, -50 ms on the x-axis indicates that we were measuring visual response strengths (y-axis value) for all trials in which stimulus onsets occurred within the interval from -75 ms to -25 ms from microsaccade onset. As can be seen, there was clear premicrosaccadic enhancement and postmicrosaccadic suppression, consistent with our earlier results ^10–12^. **(b)** Same as **a** but for negative luminance polarity stimuli. Note how there was stronger premicrosaccadic enhancement than in **a** for the 10% and 20% contrast levels (upward pointing arrows at the - 50 ms time point; see the corresponding comparison in **a**); also see Figs. 3-5. Error bars denote SEM.

Notably, relative to the no-microsaccade baseline sensitivity, premicrosaccadic enhancement was systematically stronger for the dark contrasts, especially at the moderate contrast levels of 10%-20%. Consider, for example, the 20% contrast conditions in Fig. 2. For positive luminance polarity stimuli (brights), the no-microsaccade baseline sensitivity of our population was at a normalized firing rate of 0.784 (Fig. 2a). At the center of the -50 ms time bin (that is, for stimuli occurring within the interval from -75 ms to -25 ms relative to microsaccade onset), this population firing rate was 0.866; thus, the SC population response was enhanced by 10.5% in the case of premicrosaccadic extrafoveal stimulus onsets (Fig. 2a). On the other hand, for dark stimuli of the same Weber contrast level, the no-microsaccade sensitivity was 0.754, and the premicrosaccadic sensitivity was 0.916, a 21.5% increase (Fig. 2b). Therefore, there was stronger premicrosaccadic enhancement of visual sensitivity for the negative luminance polarity condition across our population. For 10% contrasts, the premicrosaccadic enhancement for bright stimuli was 5.4% at -50 ms (Fig. 2a), whereas it was 15.2% for dark ones (Fig. 2b). Importantly, these measures were made from the very same neurons under different luminance polarity conditions. Therefore, the observations cannot be explained by arguing that different SC neurons were differentially recruited by either the positive or negative luminance polarity stimuli.

### More saturated premicrosaccadic contrast sensitivity curves for dark contrasts

Inspection of Fig. 2 in the premicrosaccadic interval also revealed that the population peak firing rates reached similar levels for 20%, 50%, and 100% contrast levels in the dark condition (Fig. 2b), but not in the bright one (Fig. 2a). This suggests that stronger premicrosaccadic enhancement for dark stimuli has led to more saturated contrast sensitivity curves. We next checked this explicitly.

We plotted the data from the time bin -50 ms (+/- 25 ms) of Fig. 2 as full contrast sensitivity curves (Fig. 3a, b). We chose this time bin because it ensured stimulus onsets before microsaccade onset, while still having close temporal proximity to the movement onset event ^10,15^ (naturally, Fig. 2 indicates that other nearby time points would have yielded similar conclusions). The black data points in Fig. 3a, b show the baseline, no-microsaccade population firing rates as a function of stimulus contrast (same values as the horizontal dashed lines in Fig. 2); the continuous curves show computational fits to a monotonically increasing mathematical function (Methods). The data and fits demonstrate expected contrast sensitivity functions of primate SC neurons ^10,38,46^. In the same panels of Fig. 3a, b, the colored data points and fits are those for only the trials in which the stimulus onset occurred within the interval from -75 ms to -25 ms relative to microsaccade onset (that is, the -50 ms x-axis point of Fig. 2). For both luminance polarities, there was clear premicrosaccadic enhancement in the population response ^10^ (Fig. 3a, b). However, at moderate contrast levels (oblique arrows), the enhancement was stronger for the negative luminance polarity stimuli (darks), as also shown in Fig. 3c, in which we plotted the difference between the premicrosaccadic and baseline contrast sensitivity curves for either dark or bright stimuli.

**Figure 3.**
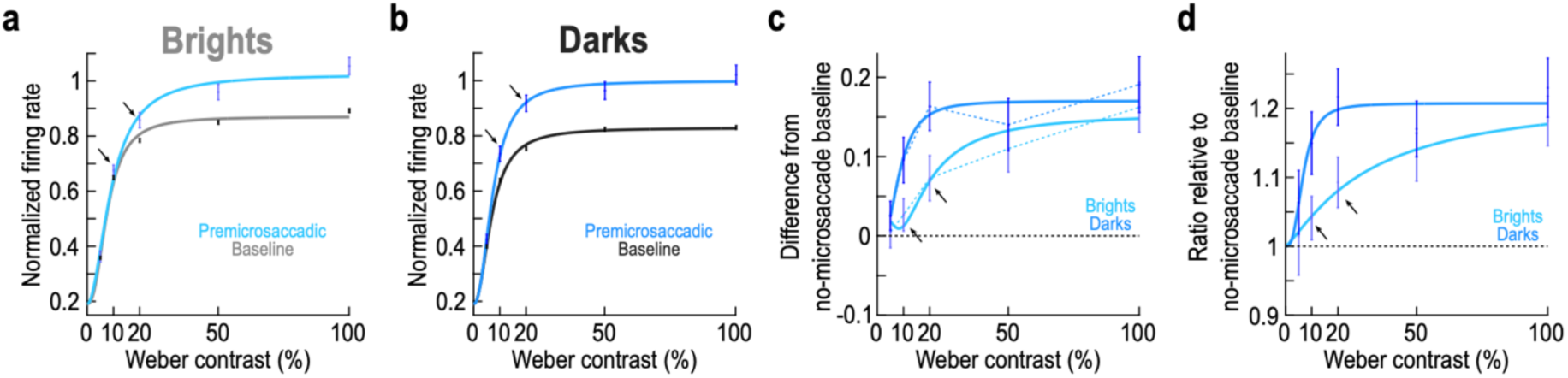
Stronger premicrosaccadic SC response gain enhancement for dark than bright stimuli at moderate contrasts. **(a)** The black data points show the normalized population visual response firing rates as a function of Weber contrast for positive luminance polarity stimuli (baseline trials). We normalized all neurons’ firing rates before averaging across them (Methods and ref. ^11^). The gray continuous curve shows a curve fit of a monotonically increasing function (Methods). In color, we show the results and curve fit for stimulus onsets within the interval from -75 ms to -25 ms relative to microsaccade onset. There was clear premicrosaccadic enhancement ^10^. However, at moderate contrast levels (e.g. black oblique arrows), the enhancement was small. **(b)** Same as **a** but for the negative luminance polarity stimuli being presented to the very same neurons. The enhancement at moderate contrast levels was now larger (e.g. black oblique arrows). **(c)** For either brights or darks, we plotted the difference between premicrosaccadic and baseline data points and curves in **a**, **b**. The difference curve for darks was systematically higher than that for brights. The black oblique arrows highlight the observation that for brights, the difference between premicrosaccadic and baseline curves was smaller than for darks at moderate contrast levels (see Table 1). Error bars denote SEM. **(d)** We also took the ratio of the premicrosaccadic to baseline data points in **a**, **b**. Here, we fitted the individual-point ratio measures to a mathematical function of the same form as that used in **a**, **b** for easier visualization (Methods). Again, at moderate contrast levels, there was stronger premicrosaccadic enhancement for dark stimuli (see Fig. 4f-j). Error bars denote SEM.

Quantitatively, at 20% contrast, premicrosaccadic enhancement for dark stimuli was already at the plateau (saturation) level of enhancement in Fig. 3c, whereas it was not at saturation for the bright stimuli (consistent with Fig. 2). Indeed, in Fig. 3a, baseline and premicrosaccadic normalized firing rates at 20% were 0.7844 (+/- 0.0278 SEM) and 0.8573 (+/- 0.0068 SEM), an enhancement of 9.293 %; on the other hand, the values in Fig. 3b were 0.7541 (+/- 0.0067 SEM) and 0.9175 (+/- 0.0298 SEM), respectively, corresponding to an enhancement of 21.668%.

To confirm these observations statistically, we performed a two-way ANOVA on the difference measures of Fig. 3c. For each neuron, we measured the visual response strength in the baseline trials and subtracted it from the visual response strength in the premicrosaccadic trials (Methods), and we did this for each contrast and luminance polarity separately. We then performed a two-way ANOVA on these difference measures, with the ANOVA factors being stimulus contrast and stimulus luminance polarity. We found a significant main effect of contrast (F=6.450085; p=3.75×10^-5^) and a significant main effect of luminance polarity (F=10.60368; p=0.001149), suggesting that premicrosaccadic enhancement (as measured by the difference between response strengths on premicrosaccadic and baseline trials) depended on both stimulus factors (contrast and luminance polarity). There was also no significant interaction between contrast and luminance polarity (F=1.781741; p=0.129839). We then performed t-tests, on the same difference measures, comparing premicrosaccadic enhancement for dark and bright luminance polarities at each contrast level. The results of these tests, after Bonferroni correction, confirmed that there was significantly stronger premicrosaccadic enhancement for dark than bright stimuli at 20% contrast (Table 1).

**Table 1.** Bonferroni-corrected t-test results between dark and bright luminance polarities at each contrast level. At each contrast level, we compared premicrosaccadic enhancement strength for dark and bright luminance polarities. The measure of premicrosaccadic enhancement in this case was obtained by subtracting baseline visual sensitivity from premicrosaccadic visual sensitivity in each condition (like visualized in Fig. 3c and the associated ANOVA). Premicrosaccadic trials were those in which the stimulus onset occurred between -75 ms and -25 ms from microsaccade onset. Baseline trials were those without any microsaccades within +/- 150 ms from stimulus onset. The shown p-values were all Bonferroni-corrected for multiple comparisons. Boldfaced p-values are those deemed significant at the 95% confidence limit (after Bonferroni correction).

|  | 5% | 10% | 20% | 50% | 100% |
| --- | --- | --- | --- | --- | --- |
| **Comparing the strength of premicrosaccadic enhancement between dark and bright luminance polarities** | p=1 | p=0.1585 | **p=0.0365** | p=1 | p=0.1865 |

We also documented the raw firing rates of the individual neurons in our population, with and without microsaccades. We did so by measuring, for each neuron, the visual response strength on baseline no-microsaccade trials and plotting it together with the visual response strength when stimulus onsets occurred within the interval from -75 ms to -25 ms relative to microsaccade onset (Methods). The results for each contrast and luminance polarity are shown in Fig. 4a-e. For each condition in the scatter plots, the x-axis value shows the baseline no-microsaccade visual response strength of a single neuron, and the y-axis value shows the response strength with premicrosaccadic stimulus onset. We then used these raw measurements to calculate the full distributions of enhancement ratios (ratios of premicrosaccadic to baseline visual response strengths) across all individual neurons (Fig. 4f-j, with the red vertical line in each histogram indicating the mean across the population). Such enhancement ratio distributions correspond to the summary population plot provided in Fig. 3d, and they provided an alternative way to assess premicrosaccadic enhancement. At 10% contrast, the mean enhancement ratio for bright contrasts was 1.015, whereas it was 1.191 for dark contrasts (Fig. 4g). Similarly, at 20% contrast, the premicrosaccadic enhancement ratio was 1.073 for brights and 1.346 for darks (Fig. 4h). These values are both qualitatively and quantitatively similar to those summarized in Fig. 3d, and they were also confirmed statistically. Specifically, we reported the Bonferroni-corrected p-values in each panel of Fig. 4f-j, obtained when comparing the ratio distributions between dark and bright stimuli at each contrast level. Once again, there was a significant effect at 20% contrast. Therefore, all results so far support our interpretation of stronger premicrosaccadic enhancement of SC visual response sensitivity for dark than bright luminance polarities, and specifically at moderate contrast levels.

**Figure 4.**
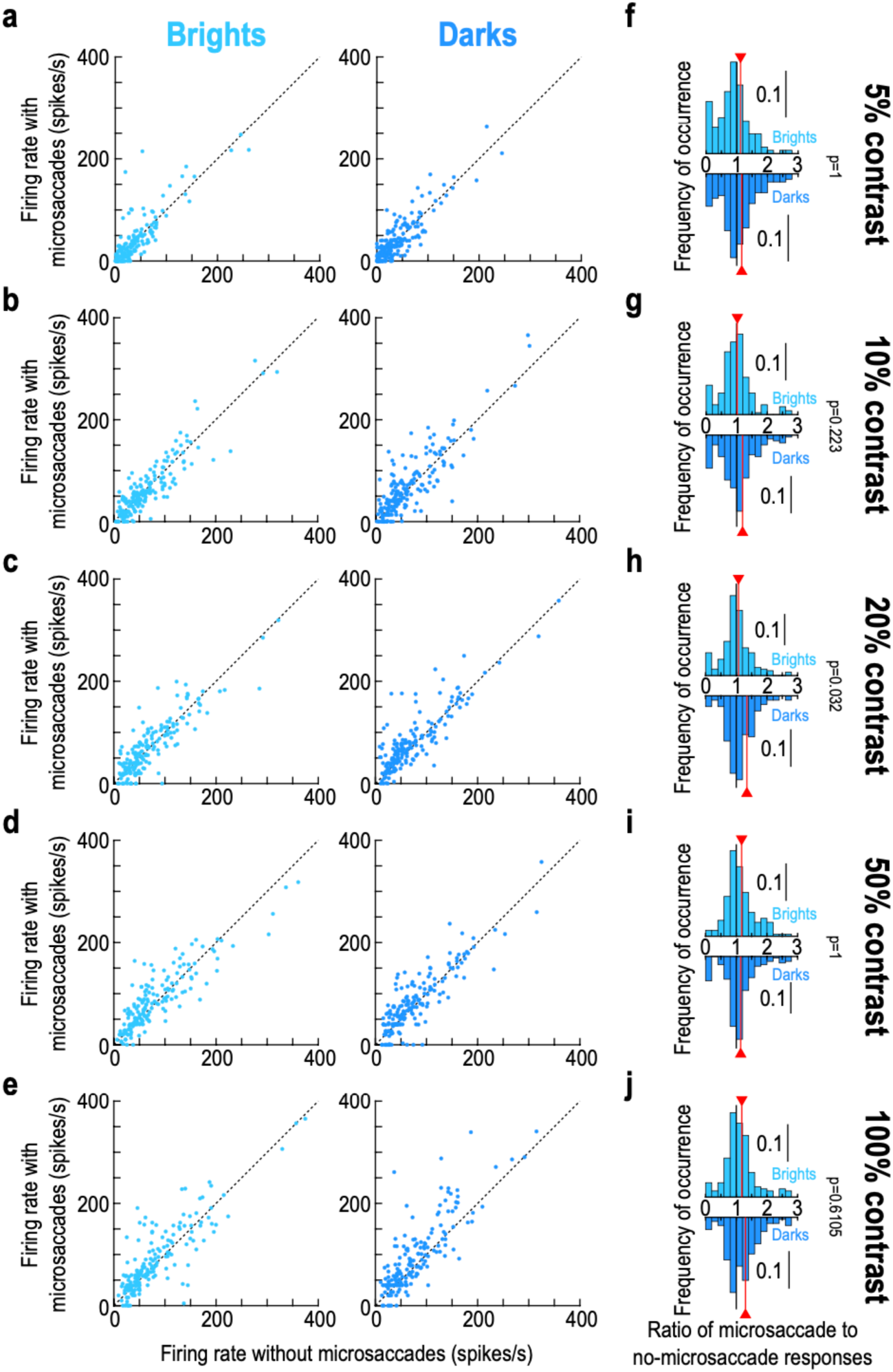
Robustness of the premicrosaccadic enhancement results at the individual neuron level. **(a)** In each panel, we plotted, for either brights (left) or darks (right), the peak visual response of neurons in baseline (x-axis) and with premicrosaccadic stimuli (y-axis; stimulus onsets within the interval from -75 ms to -25 ms relative to microsaccade onset). Each dot is an individual neuron in the 5% contrast condition. **(b-e)** At higher contrasts, premicrosaccadic enhancement was visually more robust, especially for the negative luminance polarity stimuli. **(f-j)** For each contrast level, we computed the ratio of the y- to x-axis values in **a**-**e** for each neuron and each luminance polarity. The upward pointing bar plots show the distributions of ratios for bright stimuli, and the downward pointing bar plots show the distributions of ratios for dark stimuli. In each distribution, the vertical red line indicates the mean across the population. Each p-value shows the Bonferroni-corrected result of a t-test comparing the dark to bright distribution at each contrast level. There was stronger premicrosaccadic enhancement for darks than brights at 20% contrast (see Fig. 3d). n = 166-193 neurons per panel and condition (Methods).

### Strongest enhancement levels for 20% dark contrasts

To better visualize the quantitative premicrosaccadic enhancement values across all conditions, we calculated the perimicrosaccadic modulation time courses, but now after normalizing visual responses for each condition individually (Methods). This way, regardless of the contrast or luminance polarity, and regardless of the absolute response strength to a given stimulus contrast, a normalized population firing rate of 1 would be the no-micoraccade population sensitivity for the given condition and luminance polarity. Using this approach, we obtained the time courses of Fig. 5. Each row represents one contrast level, and the two columns separate the two luminance polarities; error bars denote SEM. In each row, the leftward red arrow across the two columns indicates the peak value of the negative luminance polarity curve (in the premicrosaccadic interval), and it helps to compare the peak premicrosaccadic enhancement value to the value observed for the corresponding positive luminance polarity stimuli in the left column. As can be seen, especially for the contrast level of 20%, there was substantially stronger premicrosaccadic enhancement of SC visual neural sensitivity with dark than bright contrasts, as expected from our earlier analyses. Thus, we observed the strongest premicrosaccadic enhancement effect in the SC for 20% dark contrasts.

**Figure 5.**
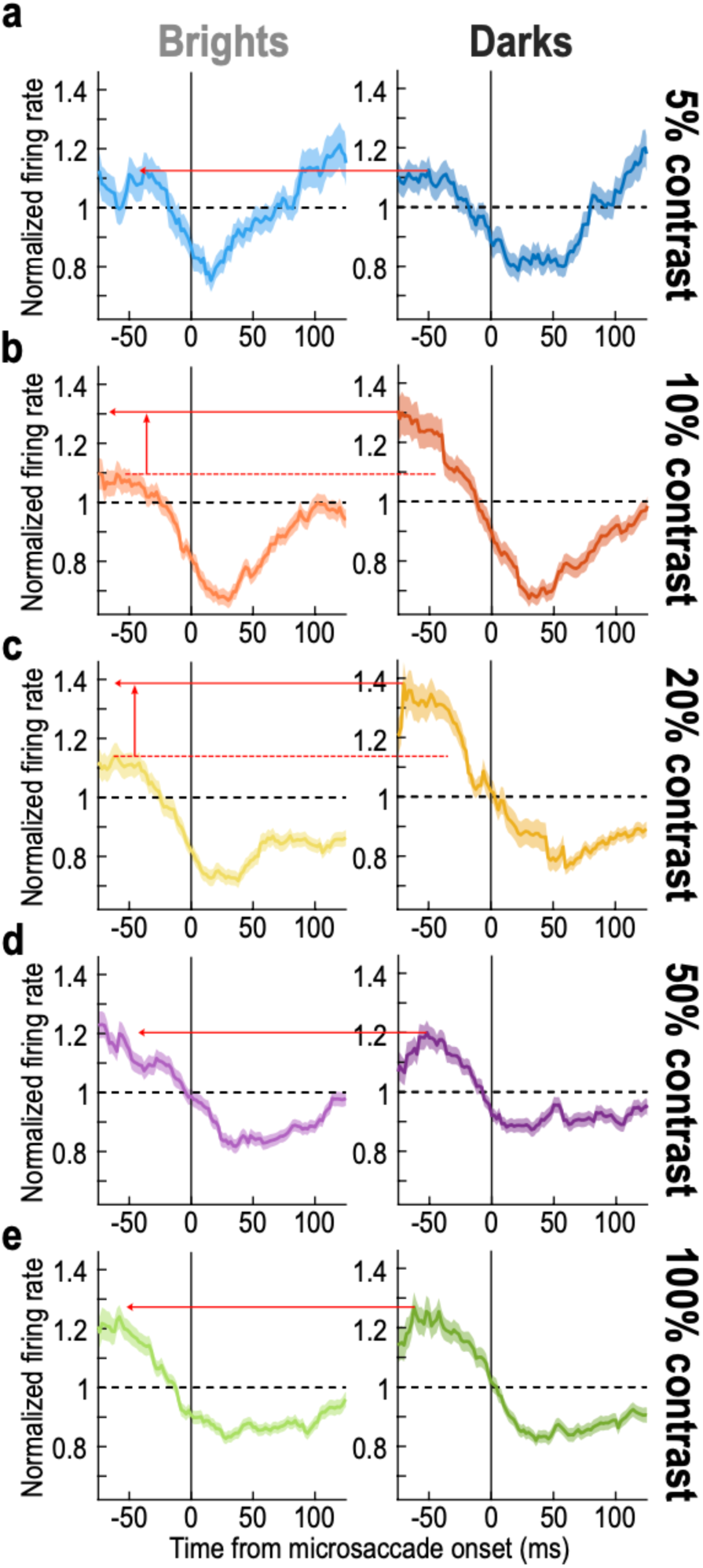
Time courses of perimicrosaccadic modulations relative to the no-microsaccade sensitivity within each contrast level and luminance polarity condition. **(a)** Each panel is similar, in principle, to what we showed in Fig. 2. However, here, all firing rates were normalized to the no-microsaccade population sensitivity of the exact condition (contrast level and luminance polarity) being analyzed by the panel. Thus, a value of 1 on the y-axis value indicates the no-microsaccade sensitivity level of the population (for a particular stimulus contrast and luminance polarity) in the absence of peristimulus microsaccades. The panel on the left shows the time course for bright stimuli, and the panel on the right shows the time course for dark stimuli. The leftward red arrow connects the peak of the curve on the right (in the prestimulus interval) to the left panel; this allows comparing the strengths of premicrosaccadic enhancement in the two panels. In this case, there was similar premicrosaccadic enhancement between dark and bright stimuli. **(b, c)** However, for 10% and 20% contrast levels, there was stronger premicrosaccadic enhancement for dark stimuli, consistent with Fig. 3. **(d, e)** This effect dissipated again for higher contrasts. Error bars in all panels denote SEM.

### Independence from individual neuron preferences for darks or brights

Because SC neurons can individually prefer either dark or bright stimuli ^38^, we additionally considered the possibility that the results above were restricted to only a subset of SC neurons (for example, those only preferring dark stimuli). We repeated the above analyses for only neurons preferring bright stimuli (126/193 neurons in our database) or only neurons preferring dark ones (67/193 neurons). We labeled the former neurons as “bright-sensitive neurons” and the latter one as “dark-sensitive neurons”.

Figure 6a, b shows the contrast sensitivity curves (similar to Fig. 3a, b) of only the bright-sensitive neurons. The baseline curves (light and dark gray colors) confirm that these neurons did indeed prefer the bright stimuli; this is because the plateau firing rates were higher for bright stimuli (Fig. 6a) than for dark ones (Fig. 6b). Interestingly, the neurons had minimal premicrosaccadic enhancement for bright stimuli of 10% and 20% contrast (black oblique arrows in Fig. 6a), but they had clear enhancement for dark stimuli of the same contrasts (black oblique arrows in Fig. 6b). Thus, the observations of Figs. 1-5 above still held for only the bright-sensitive neurons.

**Figure 6.**
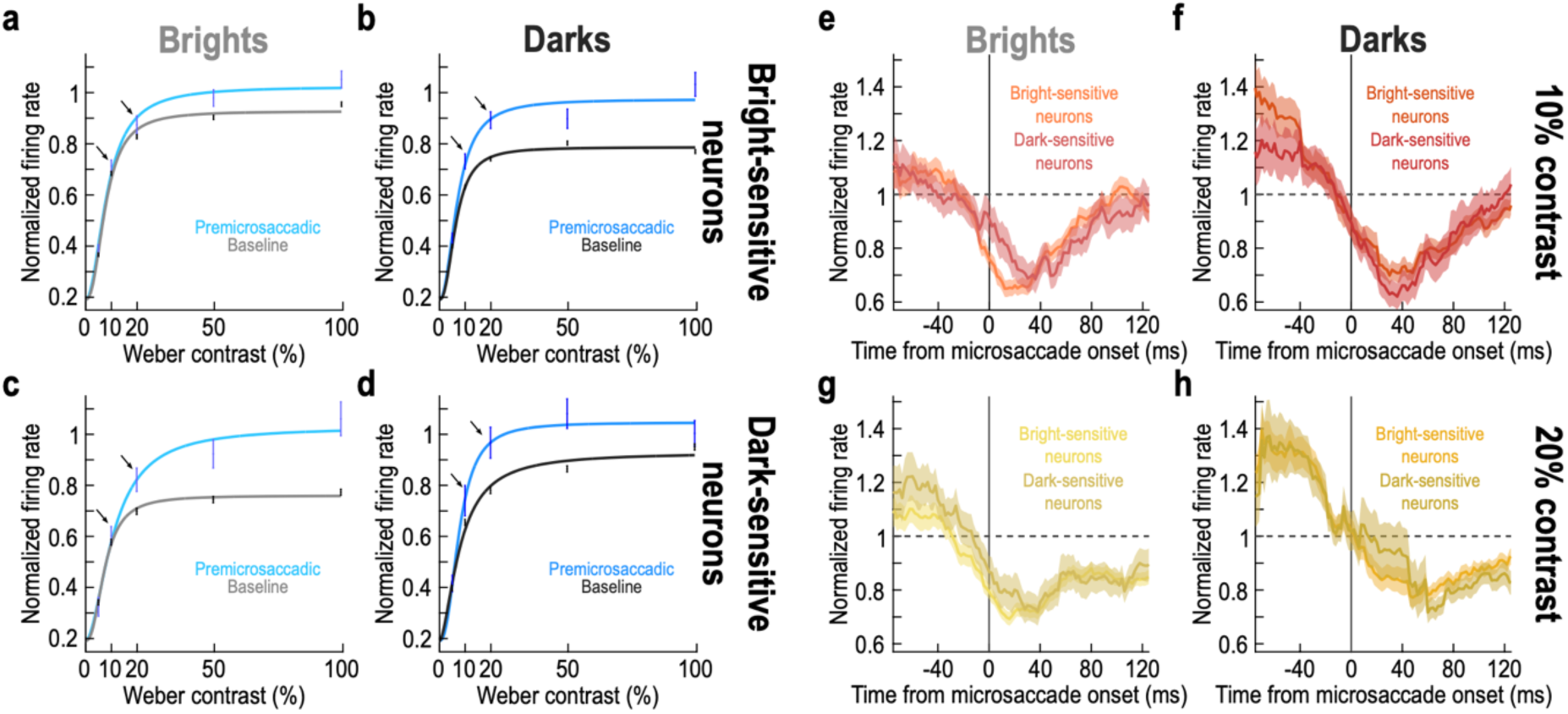
Stronger premicrosaccadic enhancement of SC visual sensitivity for moderate dark contrasts, independent of individual neuron preferences for either darks or brights. **(a, b)** Similar analyses to Fig. 3a, b, but now for only neurons preferring bright stimuli in their baseline activity. This preference can be seen by comparing the light and dark gray data points and curves across the two panels. In the premicrosaccadic interval, there was still stronger enhancement for dark stimuli at moderate contrasts (e.g. black oblique arrows). Note that there was also stronger enhancement at 100% contrast (right panel), suggesting that the preference for brights in the left panel might have caused a ceiling effect on premicrosaccadic firing rates. **(c, d)** Similar analyses for dark-sensitive neurons. Now, there was more of a ceiling effect with dark contrasts than bright ones, and this makes sense because the neurons’ baseline responses were now intrinsically stronger for darks. Nonetheless, there was still stronger premicrosaccadic enhancement for dark contrasts at moderate contrast levels (e.g. black oblique arrows). Thus, the effects of Figs. 1-5 above persisted independent of neuron preference. **(e-h)** Similar analyses to Fig. 5 but after separating neurons according to their preference for either dark or bright stimuli. Neuron preference did not alter the general observation that there was stronger premicrosaccadic enhancement for the dark stimuli of moderate contrast levels. Error bars denote SEM, and all other conventions are similar to Figs. 3, 5.

It is also interesting to note that there was generally stronger premicrosaccadic enhancement for the dark stimuli in the bright-sensitive SC neurons. For example, at 100% contrast, these neurons exhibited a sensitivity enhancement relative to baseline of only 10.15% with the preferred bright stimuli, but they showed a 34.14% premicrosaccadic enhancement for the non-preferred dark stimuli. This could reflect a ceiling effect associated with the preferred bright stimuli. In other words, the neural responses were already almost as strong as could be for the preferred stimuli (by virtue of the stimuli being preferred). As a result, there was less room for premicrosaccadic enhancement on top of the already strong visual responses. Consistent with this, in the dark-sensitive neurons instead, we now observed generally stronger premicrosaccadic enhancement for the non-preferred bright stimuli than for the preferred dark ones (Fig. 6c, d: 37.19% enhancement for 100% contrast bright stimuli but only 5.63% enhancement for the preferred 100% contrast dark stimuli).

Critically, even in the dark-sensitive neurons (Fig. 6c, d) in which premicrosaccadic enhancement was generally stronger for bright stimuli (especially at 100% contrast), there was still stronger premicrosaccadic enhancement for dark stimuli at the moderate contrast levels (e.g. black oblique arrows in Fig. 6c, d). Thus, even though the firing rates in these neurons were approaching a ceiling effect due to their preference for dark stimuli, at the moderate contrast levels, when there was still room for response strength enhancement compared to 100% contrasts, the dark-sensitive neurons still experienced stronger premicrosaccadic enhancement for dark than bright stimuli. Indeed, at the moderate contrasts, the baseline sensitivity curve of these dark-sensitive neurons was lower than the baseline sensitivity curve of bright-sensitive neurons for their preferred luminance polarity (compare the gray, baseline, curves in Fig. 6a, d at 20% contrast levels). This is further evidence that the dark-sensitive neurons had a better chance to avoid ceiling effects for dark stimuli of moderate contrast levels, and therefore still experience stronger premicrosaccadic enhancement at 20% contrast levels.

Statistically, we ran the same ANOVA as that we ran above on the main neural population (calculating the difference measure of premicrosaccadic enhancement, and related to Fig. 3c). Here, we added neuron preference as an additional factor of the ANOVA. There was a main effect of contrast (F=5.622236; p=0.00017) and luminance polarity (F=5.531184; p=0.0188), but not of neuron preference (F=0.043712; p=0.835426), suggesting that our results of Figs. 1-5 above did not depend on neuron preferences for either brights or darks. Moreover, the only significant interaction term was that between luminance polarity and neuron preference (F=5.607694; p=0.017987), consistent with the ceiling effects mentioned above (Fig. 6a-d).

Consistent with the above, full time courses of perimicrosaccadic sensitivity modulations exhibited similar trends in our bright- and dark-sensitive neurons as they did in our aggregate population. For example, Fig. 6e-h shows the time courses for 10% and 20% contrasts, in a format identical to that shown in Fig. 5. The only difference is that we now plotted the time courses separately for either the dark-sensitive or bright-sensitive SC neurons. The same trends as those observed in Fig. 5 were evident. Of course, the difference in the premicrosaccadic interval between darks and brights was quantitatively smaller for the dark-sensitive neurons (due to the potential ceiling and interaction effects mentioned above); however, the overall time courses were qualitatively the same. Thus, premicrosaccadic enhancement of extrafoveal SC visual sensitivity is indeed stronger for dark stimuli at moderate contrast levels.

Therefore, to summarize, at moderate contrast levels, stronger premicrosaccadic enhancement for dark-contrast responses in the SC is a general property, and it is not explained by the intrinsic preferences of individual neurons. However, for 100% contrast stimuli, the absolute value of premicrosaccadic enhancement for either brights or darks can depend on neuron preferences, likely reflecting ceiling effects associated with the preferred luminance polarities.

### Similar postmicrosaccadic suppression for darks and brights

Our results so far indicate that SC neurons robustly exhibit premicrosaccadic enhancement in their visual sensitivity (Figs. 1-3) ^10^, that such enhancement is specifically stronger for dark luminance polarities of moderate contrast levels (Figs. 4, 5), and that the enhancement is independent of the intrinsic preferences of individual SC neurons (Fig. 6). For the sake of completeness, and because our experiments automatically also provided us with postmicrosaccadic stimulus onsets as well, we close this study by documenting our observations on microsaccadic suppression of SC visual sensitivity.

Already in the time courses of Figs. 2, 5, 6, we could clearly see that stimulus onsets after microsaccade onset were systematically associated with suppressed SC visual sensitivity. This is consistent with our earlier observations ^10–12^, and it was additionally evident at the individual neuron level. For example, in Fig. 7a, b, we plotted the responses of the same example neuron of Fig. 1g, h, but this time for postmicrosaccadic stimuli. Microsaccadic suppression of visual sensitivity was evident.

**Figure 7.**
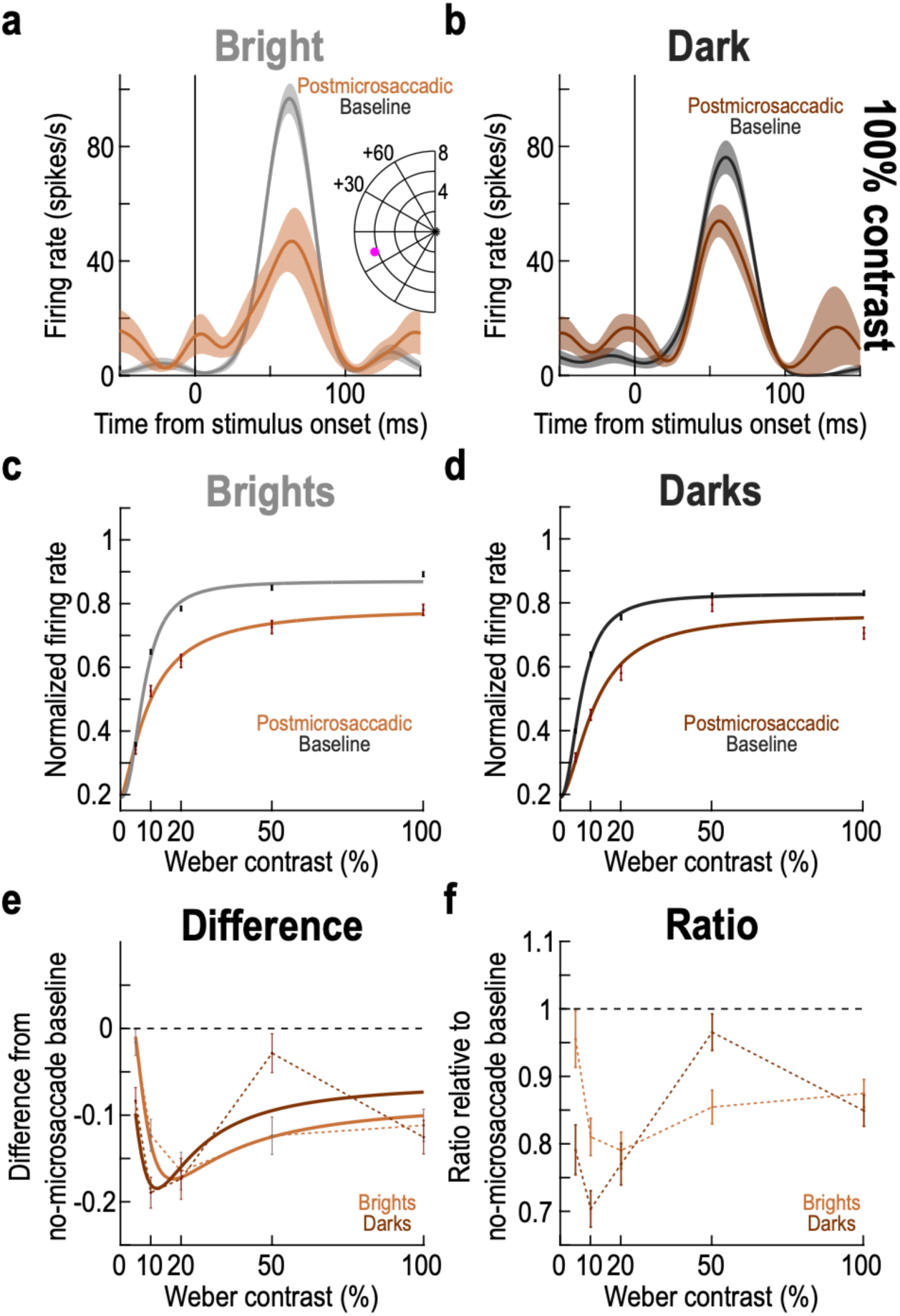
Similar postmicrosaccadic suppression of SC visual sensitivity for positive and negative luminance polarities. **(a, b)** Visual responses of the same example neuron as that shown in Fig. 1e-h. However, here, we show the responses for postmicrosaccadic stimuli instead of premicrosaccadic ones (note that only in these two panels, we included all trials with stimulus onsets within less than +100 ms from microsaccade onset, just to stay consistent with the example neuron measurement intervals of Fig. 1). As can be seen, there was expected suppression of SC visual sensitivity. n = 31-33 baseline trials and 8-10 postmicrosaccadic trials. **(c, d)** Contrast sensitivity curves like in Fig. 3a, b. Here, we compared baseline curves to those obtained with stimulus onsets occurring within the interval of +25 ms to +75 ms from microsaccade onset. There was similar suppression across luminance polarities (compare **c** and **d**). **(e, f)** Same analyses as in Fig. 3c, d, but for the postmicrosaccadic stimulus trials. At moderate contrast levels (e.g. 20%), there was similar suppression, unlike in the premicrosaccadic intervals in Figs. 1-6. The solid curves in **e** show the difference in the fitting curves, like we did in Fig. 3c. Note that in **f**, we did not apply a fit like in Fig. 3d, because the ratio measure was now not monotonic. However, the data points still show that there was no clear differential effect at moderate contrasts like in the premicrosaccadic intervals. Error bars denote SEM, and all conventions in this figure are similar to those in Figs. 1, 3.

Across the population, we repeated the analyses of Fig. 3, but this time finding stimulus onsets within the interval from +25 ms to +75 ms from microsaccade onset (Methods). These times were in the same range as the premicrosaccadic time intervals that we tested above, but they were sufficiently long after microsaccade onset to allow correctly probing postmicrosaccadic visual sensitivity. The contrast sensitivity curves of the neurons were expectedly suppressed (Fig. 7c, d) ^10^, but there was no evident difference between the suppression strengths for brights and darks. Indeed, both the difference (Fig. 7e) and ratio (Fig. 7f) curves demonstrated similar suppression effects for darks and brights, including at the moderate contrast levels (e.g. 20%). We also confirmed this statistically by repeating the two-way ANOVA on the underlying individual neuron data of Fig. 7e (just like we did in association with Fig. 3c for premicrosaccadic epochs). There was a main effect of contrast (F=7.7519787; p=3.42×10^-6^) but no main effect of luminance polarity (F=0.1081618; p=0.7422842). There was also no significant interaction between contrast and luminance polarity (F=1.113516; p=0.348386). Therefore, postmicrosaccadic suppression did not depend on luminance polarity.

We also noted in Fig. 7e, f that at 50% contrast, there seemed to be much weaker suppression for dark than bright stimuli. We are not sure why this is the case. In part, it could be due to the particular choice of time window of analysis that we used. Specifically, the time courses in Figs. 2, 5 revealed a transient increase in visual sensitivity, particularly at 50% contrast, at the +50 ms time point. Whether this is just due to data variability or a real feature of postmicrosaccadic SC visual sensitivity remains to be investigated in the future. However, what is clear is that the differential effects that we observed earlier at moderate contrast levels for premicrosaccadic enhancement were not present at the same contrasts for postmicrosaccadic suppression (this is also evident by inspecting the postmicrosaccadic intervals in the time courses presented in Fig. 5 above).

Finally, we also plotted the individual neuron results, like we did for Fig. 4. As expected, we observed robust postmicrosaccadic suppression (Fig. 8), but there was again less of a differential between darks and brights than with premicrosaccadic enhancement. For example, at 10% contrast, the mean population suppression ratio for brights and darks was 24.33% and 23.24% respectively (Fig. 8g), demonstrating similar suppression strengths across luminance polarities. Similarly, the suppression ratio was 17.12% and 21.69% for brights and darks, respectively, at 20% contrast (Fig. 8h), which was again a smaller difference between luminance polarities than in the premicrosaccadic enhancement effects of Figs. 1-6.

**Figure 8.**
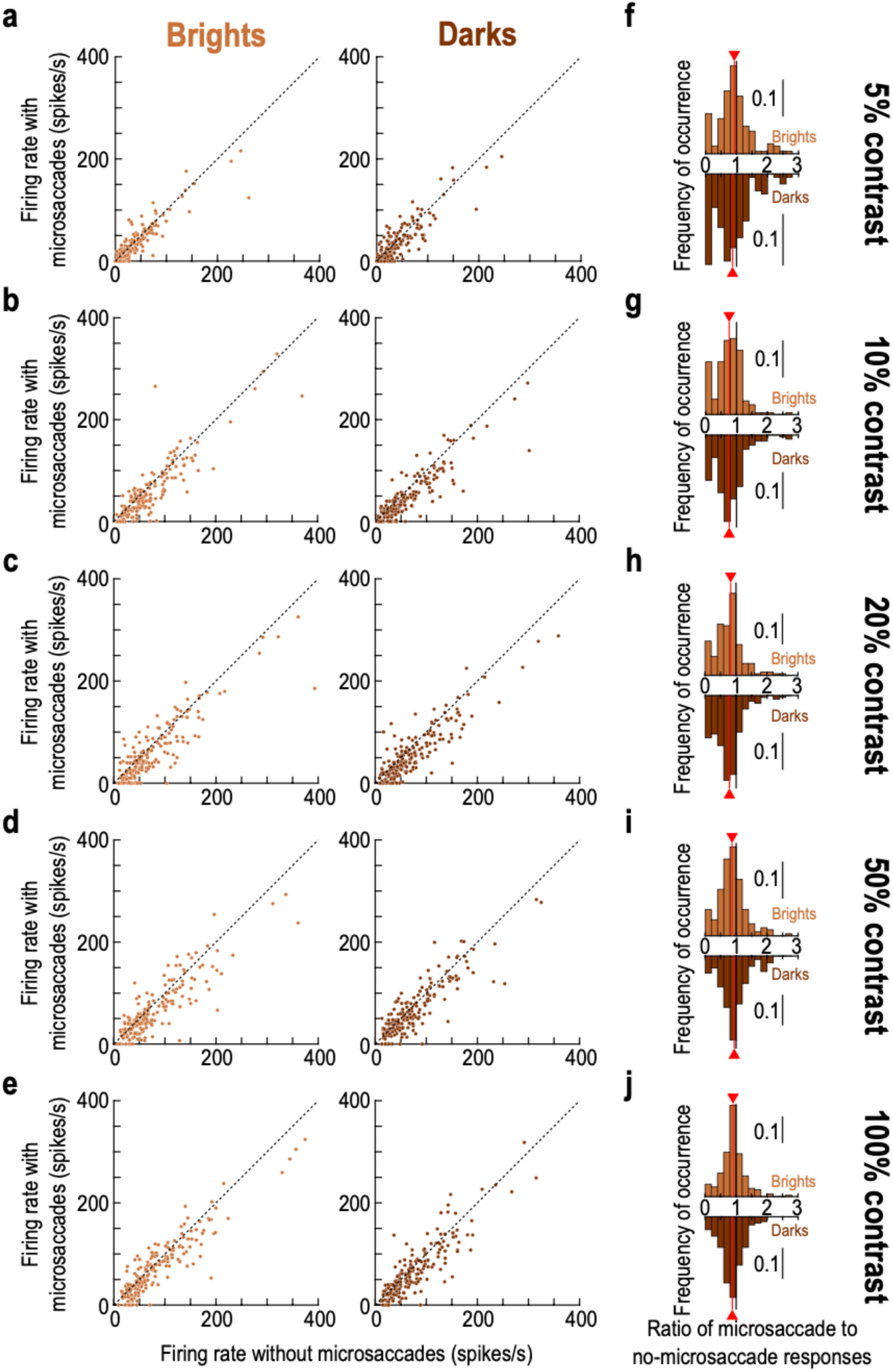
Similarity of postmicrosaccadic suppression between positive and negative luminance polarity stimuli. This figure is formatted similarly to Fig. 4. The only difference is that we now measured visual responses of individual neurons for stimulus onsets occurring within the interval from +25 ms to +75 ms from microsaccade onset. We noticed robust microsaccadic suppression of visual sensitivity. Critically, the histograms in **f**-**j** show that there were similar suppression strengths across all contrasts for the positive and negative luminance polarity stimuli (compare the red lines in each panel). Thus, the differential effect at moderate contrast levels in the premicrosaccadic interval was specific to premicrosaccadic enhancement and did not extend to postmicrosaccadic suppression. n = 175-193 neurons per condition and panel (Methods).

Therefore, postmicrosaccadic suppression of SC visual sensitivity was similar for both dark and bright stimuli. This is also consistent with our observations in human perceptual experiments that saccadic suppression strength was similar for either bright or dark luminance probes ^36,37^.

## Discussion

In this study, we investigated the dependence of premicrosaccadic visual sensitivity enhancement in the SC on luminance polarity. We found stronger premicrosaccadic enhancement for dark versus bright stimuli having moderate contrast levels. This effect persisted even when we specifically looked at either only dark-sensitive or only bright-sensitive SC neurons. On the other hand, postmicrosaccadic suppression of extrafoveal visual sensitivity was similar for both dark and bright stimuli. Our results add to a body of evidence that ON- and OFF-channel visual processing can be different in different parts of the brain’s visual processing hierarchy ^48–53^, and these results also demonstrate that such asymmetries can even appear in the oculomotor system and during the highly specific transient phase of presaccadic vision.

From a visual processing perspective, we find it intriguing to see a dependence of premicrosaccadic visual sensitivity enhancement on luminance polarity in the SC. For one, this helps reinforce the idea that visual processing in the SC is a reformatted version of its inputs ^44^, and may not simply inherit all of its properties from the primary visual cortex (V1), a major source of anatomical connection to the SC. That is, even if V1 neurons are dominated by dark-sensitive neurons ^53^, this is not necessarily the case in SC neurons ^38^. Indeed, in our current database, we had more neurons preferring bright than dark luminance polarities ^38^, suggesting that it should not be automatically assumed that luminance polarity asymmetries in the cortex would be identically observed in the SC. Our current results add that any asymmetry that may exist in the SC is further sculpted and modified specifically during the premicrosaccadic interval, and we think that this time specificity of the effect is noteworthy. We also think that this effect is unlikely to be similarly observed, whether qualitatively or quantitatively, in V1 perimicrosaccadic visual neural sensitivity.

Even in terms of the direct anatomical projections from V1 to SC, while V1 is indeed dominated by a preference for dark stimuli ^53^, this asymmetry is less prevalent in the deeper V1 layers directly projecting to the SC ^53^. Thus, it is, again, not trivial to suggest that a luminance polarity asymmetry should be identically observed in the SC, let alone in the premicrosaccadic interval. This interval is very brief (less than ∼100 ms), and it is interesting that vision (at least at the level of the SC) during such a transient phase can preferentially favor dark stimuli. At the very least, we may safely conclude that at a specific time point in relation to microsaccades, the strong alterations in vision that are already known to take place in the SC ^9–12,15^ must also include a preferential enhancement of extrafoveal sensitivity for negative luminance polarity stimuli of moderate contrast levels. It will be important in future studies to consider the functional implications of this observation. For example, there could be ecological benefits to differential processing of negative luminance polarity stimuli in general ^42,43,54,55^, and we can now begin to ask whether ecological and functional benefits may also exist in the premicrosaccadic interval as well. Indeed, we expect similar results in premovement intervals for larger saccades, and we believe that this could be a means to equalize the contrast sensitivity curves of SC neurons for the particular case of dark luminance polarities. That is, if moderate contrasts experience strong presaccadic enhancement, then this brings the contrast sensitivity curves associated with dark stimuli closer to a saturation regime for a larger range of stimulus contrast levels. This would enhance the ability of the SC to act as a robust “detector” for dark stimuli, which seem to generally be indeed ecologically relevant ^42^.

Independent of ecological considerations, we can already think of one potential functional implication of our results so far. Specifically, microsaccades are related to peripheral covert visual attention ^20,56^. Moreover, we showed previously that there could be an almost deterministic link between the occurrence of individual microsaccades and the observation of attentional performance alterations in any individual given trial ^15,20^. We also showed that this link between individual microsaccades and perceptual performance in attentional cueing tasks is specifically mediated by premicrosaccadic enhancement of visual sensitivity in different sensory-motor brain areas ^9,10^, and that premicrosaccadic enhancement (including in perception ^15^) is actually entirely sufficient (at least theoretically) to explain classic peripheral attentional cueing effects ^15,16^. If that is the case, then our current results suggest that one can further enhance attentional cueing effects in covert visual attention experiments by using negative luminance polarity perceptual discrimination stimuli of intermediate contrast. To our knowledge, classic attentional studies generally either use Gabor-like gratings ^57^ or bright targets ^15,57–59^ instead.

Irrespective of covert visual attention, our results also add to the increasing evidence that microsaccades can have a long-ranging effect on extrafoveal vision ^7^. That is, eccentricities approximately one or two orders of magnitude larger than individual microsaccade amplitudes are significantly affected by microsaccade occurrence ^9^, and it would be important to understand how this happens. One possible mechanism underlying this is lateral interactions in the SC. For example, there are indications that different layers of the SC can exhibit different amounts of lateral excitation and lateral inhibition across the SC topographic map ^60–62^. Intriguingly, in the deeper layers, there may be long-range excitation effects ^61^, which, coupled with the large foveal magnification of neural tissue that is present in the SC topographic map ^14^, could potentially mediate an excitatory influence of microsaccades on extrafoveal SC visual sensitivity. Having said that, our present work is still correlative. That is, it could be the case that it is the enhanced extrafoveal visual sensitivity itself (for whatever reason) that gives rise to a microsaccade. Thus, in our more recent followup work ^63^, it was important for us to causally test this. We created an experimental condition that exclusively influenced microsaccade generation in the absence of peripheral stimulation, and we then investigated what happens in the peripheral SC representation as a result of this. The experimental condition itself was to control foveal motor error, since it can very potently alter microsaccade directions in monkeys ^16,64^. Doing this, we could support the hypothesis that the direction of causality can indeed flow from the act of microsaccade generation itself to a concomitant influence on peripheral visual sensitivity ^63^.

At the other end of the spectrum, we observed postmicrosaccadic suppression. This is something that we routinely observe in the SC ^10–12^. Nonetheless, we find it surprising that there was no clear difference in suppression strength between bright and dark stimuli. In fact, we recently found that in the retinal origin of saccadic suppression ^36^, there is a highly substantial difference in the suppression of ON- and OFF-channel retinal circuits ^65^. While this observation could reflect species differences, we think that it may additionally be consistent with the idea that SC visual responses are distinctly reformatted versions of visual responses in other early visual areas ^44^. To test this, one needs to repeat the current experiments, but in V1 (and ideally using the very same animals that are used for the SC recordings). The prediction is that there should be different patterns of saccadic suppression in V1 between bright and dark luminance polarity stimuli. Demonstrating this would support our intuition that SC visual responses, and how they are affected by saccade generation, are not merely inherited from visual responses in early cortical visual areas ^44^.

## Methods

### Experimental animals and ethical approvals

We analyzed neural data from the same database described recently ^38^. Perimicrosaccadic modulations in that database were not documented.

Briefly, the experiments involved recording SC visual responses from two (M and A) adult, male rhesus macaque monkeys (*macaca mulatta*) fixating a small spot.

All experiments were approved by ethics committees at the regional governmental offices of the city of Tübingen (Regierungspräsidium Tübingen) ^38^, and they were in accordance with the ARRIVE guidelines. All methods were performed in accordance with the relevant guidelines and regulations. The animals are still being used in the laboratory for other projects.

### Laboratory setup and animal procedures

The monkeys were prepared previously for behavioral and neurophysiological experiments in our laboratory ^38,64,66–68^, and we tracked eye movements using the magnetic induction technique ^69,70^. The laboratory setup was described in detail earlier ^38^. Briefly, the monkeys sat comfortably in front of a display having a gray background. They fixated a fixation spot, and they received fluid reward for maintaining gaze fixation on the spot until trial end. We recorded SC activity using single- or multi-electrode arrays inserted acutely into the SC and removed after the experiments ^38^.

### Experimental procedures

The original main task of the earlier publication ^38^ was a gaze fixation task. The monkeys initially fixated a small, black fixation spot. After several hundred milliseconds, a disc of 0.51 deg radius appeared at the recorded SC response field (RF) location, and it remained on for at least 500 ms. The monkeys were trained to maintain fixation on the fixation spot, regardless of the eccentric disc appearance. Across trials, the disc could have a Weber contrast of 5%, 10%, 20%, 50%, or 100%, with Weber contrast being defined as 100*|*I_s_*-*I_b_*|/*I_b_* (*I_s_* refers to the disc’s luminance value, and *I_b_* refers to the display background’s luminance value). Moreover, the disc could either be brighter than the background (positive luminance polarity) or darker (negative luminance polarity). In the previous publication ^38^, we only accepted trials into the analysis that did not have any microsaccades occurring near stimulus onset. Here, we specifically analyzed these trials, and we used the trials without nearby microsaccades at stimulus onset as the baseline trials (see below).

### Data analysis

Microsaccades were detected in the previous publication ^38^ using our previously established criteria ^71,72^.

For every neuron, trials in which there were no microsaccades within less than +/- 150 ms from stimulus onset were considered to be our baseline trials. These were expected to be the trials in which visual responses were least affected by either past or upcoming microsaccades ^10,11^.

In the current study, we only considered neurons in which stimulus onset resulted in an elevation of a neuron’s firing rate in any of our stimulus conditions. Therefore, we excluded the few suppressed neurons from the original study. This gave us a database of 193 neurons. We considered all visually-responsive neurons because our previous work showed that premicrosaccadic enhancement occurs for either visual-only or visual-motor SC neurons ^10^.

To assess the visual sensitivity of any given SC neuron, we measured the peak stimulus-evoked response. We did this by searching, in every trial, for the maximum value of the firing rate curve of the neuron within the interval from +10 ms to +100 ms relative to stimulus onset. This interval was large enough to allow for finding the peak response even when this response was slightly delayed for the lower contrasts ^38^. We then averaged across trial repetitions of a given condition. For the baseline (no-microsaccade) response, we measured the peak visual response from only the trials without any nearby microsaccades to stimulus onset (within +/- 150 ms). To identify microsaccade-related enhancement or suppression, we measured the peak firing rate (from the same measurement interval relative to stimulus onset), but now for only subsets of trials according to the timing relationship between stimulus onset and the nearest microsaccade onset. For example, to measure premicrosaccadic enhancement, we picked only the trials in which stimulus onset occurred within the interval from -75 ms to -25 ms relative to microsaccade onset. We then found the peak, stimulus-evoked visual response strength from only these trials. If it was larger than the baseline measurement, then the neuron experienced premicrosaccadic enhancement in its visual sensitivity ^10^ (see Fig. 1 for relevant examples demonstrating the logic of this approach).

Because perimicrosaccadic modulations have distinct time courses ^10–12^, in various analyses, we repeated the above approach, but this time with different time ranges of stimulus onsets relative to microsaccade onsets. In particular, for time course analyses like in Fig. 2, we used a moving time window approach. In each time sample, we only took trials in which flash onset occurred within +/- 25 ms from the chosen time sample. Then, we measured the peak visual response on those trials. To obtain full time courses, we moved the time sample in steps of 2 ms.

Because the above time window analysis is data intensive (only small subsets of trials are included in each sampling window), we combined measurements from across neurons in our population analyses, in order to increase robustness of the analyses. To do so, we first normalized measurements from each neuron ^11^. Specifically, we divided the firing rate of every trial of every neuron by the neuron’s baseline, no-microsaccade, peak visual response measurement. In one approach (such as in Figs. 2, 3), we normalized by the peak response of the neuron from the condition causing the strongest visual burst by the neuron. For example, if a neuron had the strongest baseline visual response for 100% dark stimuli, then we divided the neuron’s firing rates in all trials (regardless of condition) by the response for 100% dark stimuli. The advantage of this is that we maintained the relative relationships of response strengths across contrasts, which in turn allowed us to obtain full contrast sensitivity curves (such as in Fig. 3). In the other approach (such as in Fig. 5), we normalized each condition individually. For example, when analyzing enhancement and suppression for 10% contrast dark stimuli, the normalization factor was the baseline, no-microsaccade, peak visual burst measurement from only trials with 10% contrast dark stimuli. After normalizing each neuron’s firing rates, we then combined trials from across all neurons ^11^.

To visualize contrast sensitivity curves, we fit the five contrast measurements of each luminance polarity to a continuous monotonic function of the form

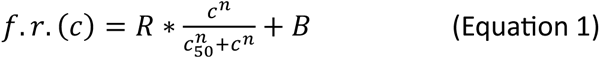

where *c* is stimulus contrast and *B*, *R*, *n*, and *c_50_* are the fit parameters ^10^. We obtained fits either for the baseline trials, or for trials in which stimulus onset occurred within a given time window relative to microsaccade onset.

When visualizing premicrosaccadic contrast sensitivity curves in relation to the baseline curves, we also took the difference between the two curves, as well as their associated data samples (separately for each luminance polarity; Fig. 3c). The continuous curves in Fig. 3c represented the difference between the continuous curves obtained from the fitting equation above.

We also took the ratio between the premicrosaccadic and baseline visual sensitivity measurements at each contrast (Fig. 3d). We then visualized the ratio for each luminance polarity by fitting the ratio measurements using the same fitting equation above.

For checking the impacts of neuron preferences, we defined dark-sensitive neurons as those in which the condition causing the highest visual response strength in baseline trials was from a dark contrast. For example, if a neuron had the highest visual response for 100% dark contrasts, then we classified it as being dark-sensitive. Similarly, we defined bright-sensitive neurons as those in which the condition causing the highest visual response in baseline trials was from a bright contrast.

We also plotted raw firing rates premicrosaccadically and in the baseline (no-microsaccade) data. We did this at each contrast level and for each luminance polarity (Fig. 4). This, in turn, allowed us to plot distributions of the raw ratio measurements (Fig. 4). Each ratio was the visual response of a given neuron on premicrosaccadic trials divided by the visual response of the same neuron on baseline trials. If the ratio was larger than 1, then the neuron experienced premicrosaccadic enhancement.

Statistically, we checked for an effect of premicrosaccadic enhancement and luminance polarity on our neurons using ANOVA’s. In one approach, we measured, for each neuron, the peak visual response on premicrosaccadic trials (trials with stimulus onset within the interval from -75 ms to -25 ms relative to stimulus onset), and we subtracted from it the peak visual response on baseline trials from the very same condition (contrast and luminance polarity). This way, all measurements within a given contrast were relative to the baseline visual sensitivity of that contrast. Then, we input these difference measurements of all neurons into a two-way ANOVA with stimulus contrast and stimulus luminance polarity as the ANOVA factors. We then ran t-tests comparing the difference measure for dark luminance polarities to the difference measure for bright luminance polarities at each contrast level individually, and we applied a Bonferroni correction to the p-values reported in Table 1.

In a second approach, we took the ratio of premicrosaccadic to baseline visual response strength. This gave the distributions in Fig. 4f-j. For each contrast level, we had one distribution of ratios for dark stimuli and one for bright stimuli. We compared the two distributions and reported the Bonferroni-corrected p-values in Fig. 4f-j.

When analyzing the influence of neuron preference, we ran the same ANOVA as that described above, but we converted it to a three-way ANOVA by adding neuron preference (dark-sensitive or bright-sensitive) as the third ANOVA factor.

Finally, we repeated all of the above analyses for trials in which stimulus onset occurred postmicrosaccadically (Figs. 7, 8). The only difference was the choice of interval dictating which trials to include. Specifically, for the statistical tests and for the summary figures not showing full time courses, we included trials in which stimulus onset occurred from +25 ms to +75 ms relative to microsaccade onset. This allowed us to assess postmicrosaccadic suppression strength, rather than premicrosaccadic enhancement.

## Acknowledgements

We were funded by the Deutsche Forschungsgemeinschaft (DFG; German Research Foundation) through the Special Priority Programme: SPP 2205 Evolutionary Optimization of Neuronal Processing (project number: HA 6749/3-2).

## Author contributions

W.W. and Z.M.H. contributed to the conception and design of the study. W.W. and Z.M.H. analyzed the data. W.W. and Z.M.H. interpreted the results. W.W. and Z.M.H. wrote the manuscript.

## Data availability statement

The datasets generated during and/or analyzed during the current study are available from the corresponding author on reasonable request.

